# DNA methylation age analysis of rapamycin in common marmosets

**DOI:** 10.1101/2020.11.21.392779

**Authors:** Steve Horvath, Joseph A. Zoller, Amin Haghani, Ake T. Lu, Ken Raj, Anna J. Jasinska, Julie A. Mattison, Adam B. Salmon

## Abstract

Human DNA methylation data have previously been used to develop highly accurate biomarkers of aging (“epigenetic clocks”). Subsequent studies demonstrate that similar epigenetic clocks can also be developed for mice and many other mammals. Here, we describe epigenetic clocks for common marmosets (*Callithrix jacchus*) based on novel DNA methylation data generated from highly conserved mammalian CpGs that were profiled using a custom Infinium array (HorvathMammalMethylChip40). From these, we developed and present here, two epigenetic clocks for marmosets that are applicable to whole blood samples. We find that the human-marmoset clock for relative age exhibits moderately high age correlations in two other non-human primate species: vervet monkeys and rhesus macaques. In a separate cohort of marmosets, we tested whether intervention with rapamycin, a drug shown to extend lifespan in mice, would alter the epigenetic age of marmosets, as measured by the marmoset epigenetic clocks. These clocks did not detect significant effects of rapamycin on the epigenetic age of marmoset blood. The common marmoset stands out from other mammals in that it is not possible to build accurate estimators of sex based on DNA methylation data: the accuracy of a random forest predictor of sex (66%) was substantially lower than that observed for other mammals (which is close to 100%). Overall, the epigenetic clocks developed here for the common marmoset are expected to be useful for age estimation of wild-born animals and for anti-aging studies in this species.

## Introduction

Substantial progress in the basic biology of aging, including discovery of interventions that extend longevity, have come from studies of invertebrates and rodents. It remains challenging to translate this basic research to clinical application in relatively healthy and genetically varied groups of people. Rodents such as mice and rats are only distantly related to humans and have undergone different evolutionary pressures that may have driven species-specific idiosyncrasies of aging. Due to the long lifespans of humans, any outcomes of longevity interventions in human studies are likely to require very long follow up at high cost. To overcome this hurdle, a case can be made for testing promising interventions in non-human primate (NHP) models. The long-term calorie restriction studies utilizing rhesus macaques (*Macaca mulatta*) are a prime example of this approach of using NHP for studying aging interventions. While rhesus macaques are attractive as a model of primate aging due to its high DNA sequence homology with humans (around 92%), the average lifespan of this species in captivity is around 27 years (maximal lifespan is 42 years)^1,2^. The long lifespans of rhesus monkeys required decades-long commitment to accomplish the goals of the calorie restriction study ^3–6^.

Among the common NHP species maintained for research purposes, the common marmoset (*Callithrix jacchus*), a small NHP native to South America, has one of the shortest natural lifespans among anthropoid primates with an average lifespan in captivity of approximately 7–8 years and maximum lifespans reported between 16 and 21 years^7,8^. Further, marmosets that live in a closed colony have a natural adult mortality that drives a decline in their cumulative survival rate from about 85 to 35% that generally occurs between 5 and 10 years of age^7^. Thus, the significantly shorter lifespan of marmosets has some advantages over other longer-lived NHP or humans in terms of developing and managing studies on the aging process^9^. In particular, the relatively short lifespan of marmosets is particularly attractive for longitudinal studies addressing the impact of interventions on aging since such data on lifespan from such studies are likely to be available more quickly than if performed using other NHP.

Due to the growing interest in using marmosets for aging research, it is of interest to develop and refine tools that can complement existing functional and physiological geroscience research in this species. Such tools will impact both our understanding of the biology of aging in this species (and its context with other species) as well as help determine, in particular, whether anti-aging interventions that affect biomarkers of aging in one species also slow corresponding biomarkers in marmosets. This article focuses on the development of DNA methylation-based biomarkers (epigenetic clocks) for blood samples from marmosets.

With the technical advancement of methylation array platforms that can provide quantitative and accurate profiles of specific CpG methylations, came the insight to combine methylation levels of several DNA loci to develop an accurate age estimator (reviewed in ^10,11^). For example, the human multi-tissue epigenetic age estimation method combines the weighted average of methylation levels of 353 CpGs into an age estimate that is referred to as DNAm age or epigenetic age ^12^. Most importantly, we and others have shown that human epigenetic age relates not just to chronological age, but biological age as well. This is demonstrated by the finding that the discrepancy between DNAm age and chronological age (what we term “epigenetic age acceleration”) is predictive of all-cause mortality in humans even after adjusting for a variety of known risk factors ^13–17^. Moreover, we have demonstrated that the human pan-tissue epigenetic clock applies to chimpanzees^12^ without change, but it loses utility for other animal species likely as a result of evolutionary genome sequence divergence from humans. Recently, several groups have tested whether gold standard longevity interventions in mice also affect epigenetic clocks in this species ^18–23^. Overall, these independent observations led to the important insight that interventions such as calorie restriction and reduction of growth hormone signaling significantly reduce biological age as measured by these epigenetic clocks.

Our current study addressed several aims: First, we developed a DNA-methylation based estimator of chronological age across the entire lifespan of common marmosets. Second, we used this novel biomarker to determine the effects of a putative anti-aging intervention (rapamycin) in marmosets. Third, we evaluated the extent to which marmoset-specific biomarkers of aging apply to Old World NHP (vervet monkey, rhesus macaque) and to humans. Fourth, we characterized individual CpGs that relate to chronological age and rapamycin treatment in marmosets.

## Results

We generated high quality DNA methylation data from n=96 blood samples taken from marmosets of ages across a large proportion of the lifespan of this species (approximately 0.5 – 15.5 y of age, **Table 1**). A putative outlier (possibly platemap error) was removed resulting in N=95 blood samples. The blood samples came from 2 groups of animals: those that were used for developing an epigenetic clock (training set) and those that were from an interventional study, testing the effect of rapamycin (test set). The treated animals had a mean age of 10 at the time of the blood draw (**Table 1**). Unsupervised hierarchical clustering reveals that the samples largely clustered by tissue type (**Supplementary Figure 1**).

**Table 1.**
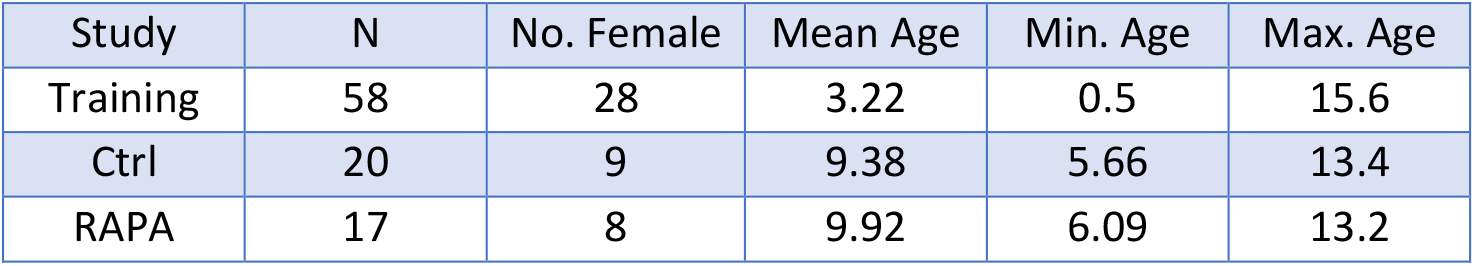
Description of the DNA methylation data. Study denotes the training set for building the epigenetic clocks. RAPA=treated with rapamycin. Ctrl=control group. N=Total number of blood samples per study. Number of females. Age (mean, minimum and maximum).

| Study | N | No. Female | Mean Age | Min. Age | Max. Age |
| --- | --- | --- | --- | --- | --- |
| Training | 58 | 28 | 3.22 | 0.5 | 15.6 |
| Ctrl | 20 | 9 | 9.38 | 5.66 | 13.4 |
| RAPA | 17 | 8 | 9.92 | 6.09 | 13.2 |

### Epigenetic clocks

Our different clocks for marmoset can be distinguished along two dimensions: species and measure of age. The marmoset pan-tissue clock was trained to apply only to blood samples. The two human-marmoset dual-species epigenetic clocks are distinct, by way of measurement parameters. One estimates *chronological age* (in units of years), while the other estimates *relative* age, which is the ratio of chronological age to maximum lifespan; with values between 0 and 1. This ratio allows alignment and biologically meaningful comparison between species with very different lifespan (marmoset and human), which is not afforded by mere measurement of chronological age.

To arrive at unbiased estimates of the epigenetic clocks, we used cross-validation of the training data. The cross-validation study reports unbiased estimates of the age correlation R (defined as Pearson correlation between the age estimate (DNAm age) and chronological age as well as the median absolute error. As indicated by its name, the marmoset blood tissue clock is highly accurate in age estimation of the donors of blood samples (R=0.95 and median absolute error 0.72 years, **Figure 1A**). The human-marmoset clock for relative age is highly accurate when both species are analyzed together (R=0.96, **Figure 1B**) but lower when the analysis is restricted to marmoset blood samples (R=0.86, **Figure 1C**). This demonstrates that relative age circumvents the skewing that is inherent when chronological age of species with very different lifespans are measured using a single formula.

**Figure 1:**
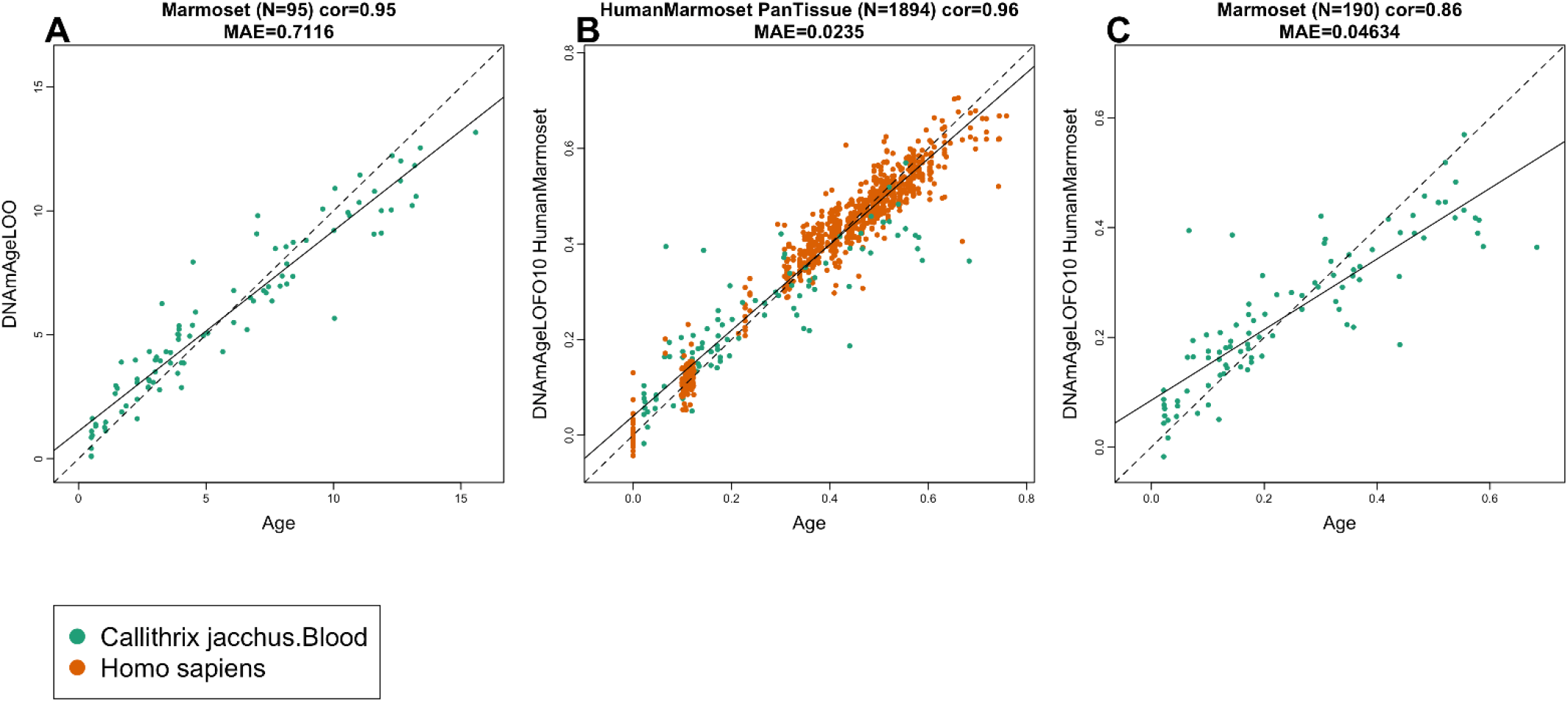
Cross-validation study of epigenetic clocks for common marmosets and humans. A) Epigenetic clock for blood samples from marmoset. Leave-one-sample-out estimate of DNA methylation age (y-axis, in units of years) versus chronological age. B) Ten fold cross validation analysis of the human-marmoset clock for relative age. Dots are colored by tissue type (green=marmoset). C) Excerpt from panel B but restricted to marmosets. Each panel reports the sample size, correlation coefficient, median absolute error (MAE).

### Application to other primates

To determine the cross-tissue performance and the cross-species conservation of the marmoset clocks, we applied the marmoset clocks two Old World primate species, vervet monkeys (n=240 from 3 tissues) and rhesus macaque (n=283 samples from eight tissues). The data from vervet monkeys (*Chlorocebus sabaeus*) and rhesus macaques are described in companion papers ^24,25^.

The marmoset clocks exhibit moderately high age correlations in vervet monkeys (R=0.8, **Figure 2A**). The human-marmoset clock for chronological age and relative age lead to similar age correlations (R=0.89 and 0.88, respectively, **Figure 2B,C**). By comparison, the marmoset clocks are less accurate in rhesus macaques (Figure 3): pure marmoset clock (r=0.4, **Figure 3A**), human-marmoset clocks for chronological and relative age (R=0.63 and 0.73, **Figure 3B,C**).

**Figure 2.**
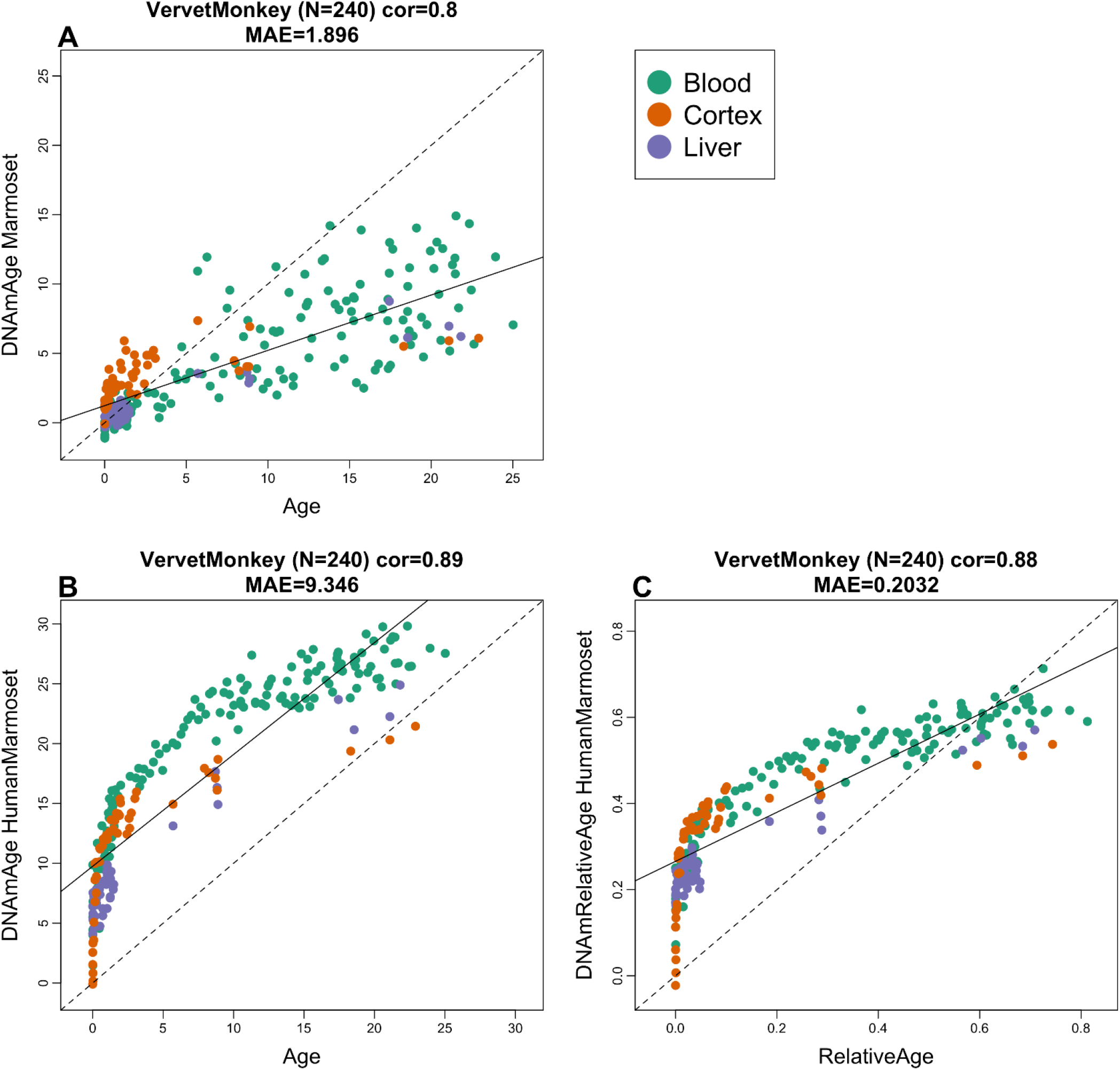
Marmoset clocks applied to tissues from vervet monkey. Each dot corresponds to a tissue sample from vervet monkey (Chlorocebus sabaeus). Chronological age of the vervet specimens (x-axis) versus DNAm age estimate of the marmoset A) blood clock, B) human-marmoset clock for chronological age, E) human-marmoset clock for relative age. The number of samples is shown in parentheses; cor – Pearson’s correlation, MAE – median absolute error.

**Figure 3.**
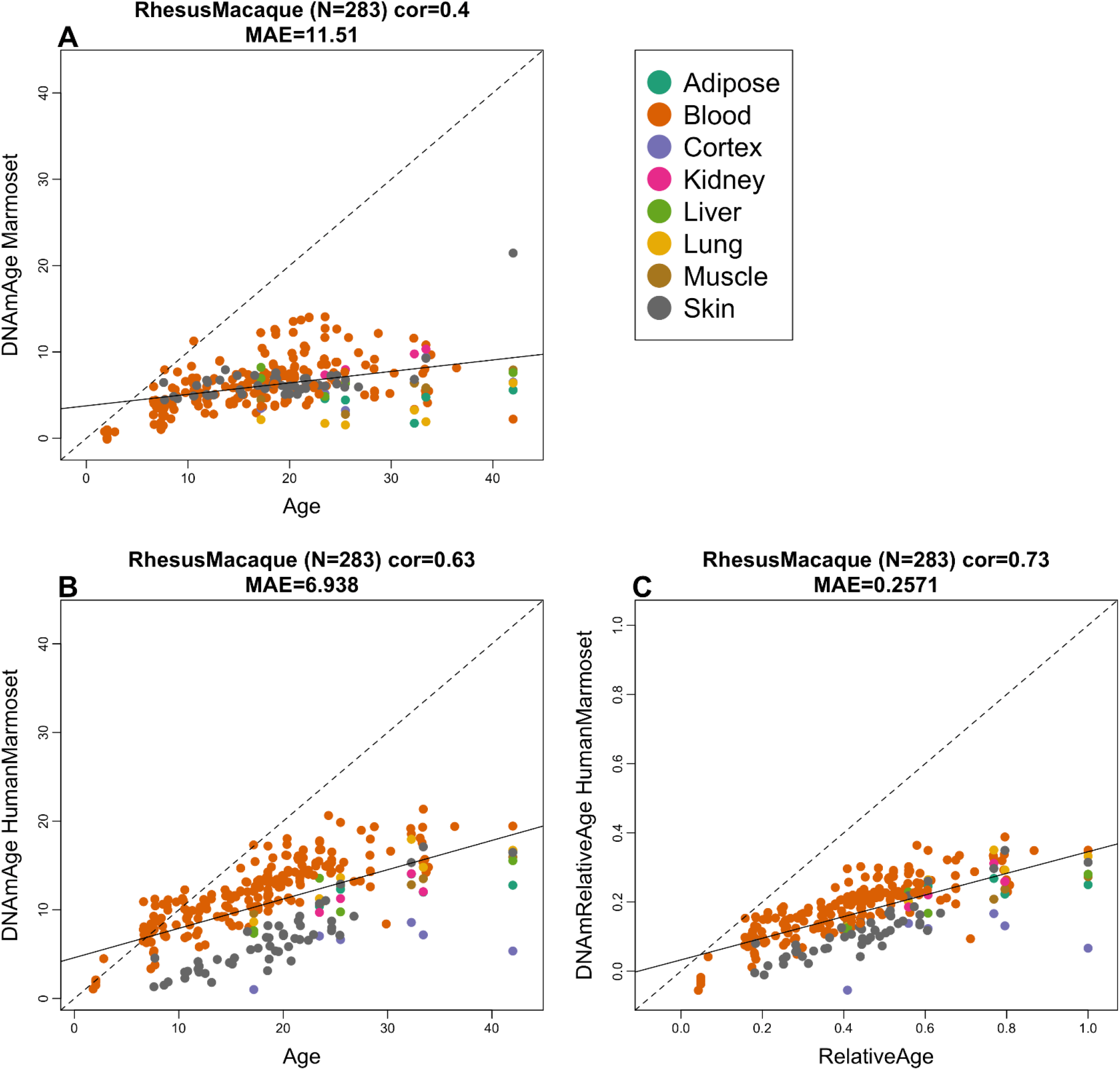
Marmoset clocks applied to tissues from rhesus macaques. Each dot corresponds to a tissue sample from rhesus macaques: Chronological age of the macaque specimens (x-axis) versus DNAm age estimate of the marmoset A) blood clock, B) human-marmoset clock for chronological age, C) human-marmoset clock for relative age. The number of samples is shown in parentheses; cor – Pearson’s correlation, MAE – median absolute error.

Overall, we find that the human-marmoset clock for relative age leads to the highest age correlations in the two non-human primates. While the human-marmoset clock for chronological age is poorly calibrated in other non-human primates, its high age correlations show that it would lend itself for ranking samples from other primates according to age (from youngest to oldest).

### Age-related and sex related CpGs

Epigenome-wide association (EWAS) of chronological age in blood from marmosets can be found in Figure 4. In general, chronological age had a larger effect size on DNAm changes with a minimum p value of 10^−25^ (**Figure 4A**). The top 1000 DNAm changes with age had a genome-wide p value of <10^−8^ significance. Hypermethylation in *ANK1* exon, *SCG3* promoter, and *UNC79* exon were the top aging-associated DNAm changes in marmoset blood. In general, most of the CpGs in the promoter and 5’UTR regions were hypermethylated with age (**Figure 4B**). This corroborate the abundance of CpG islands in the promoters and gradual hypermethylation with age in humans, rodents, and other species. The low number of sex related CpGs (**Figure 4C**) echoes our finding that one cannot build an accurate random forest predictor of sex. A transcription factor analysis of age related CpGs (**Figure 4D**) revealed only a single motif: Nuclear Transcription Factor Y Subunit Alpha (NFYA), which is involved in cancer ^26^, and Rheumatoid arthritis ^27^.

**Figure 4.**
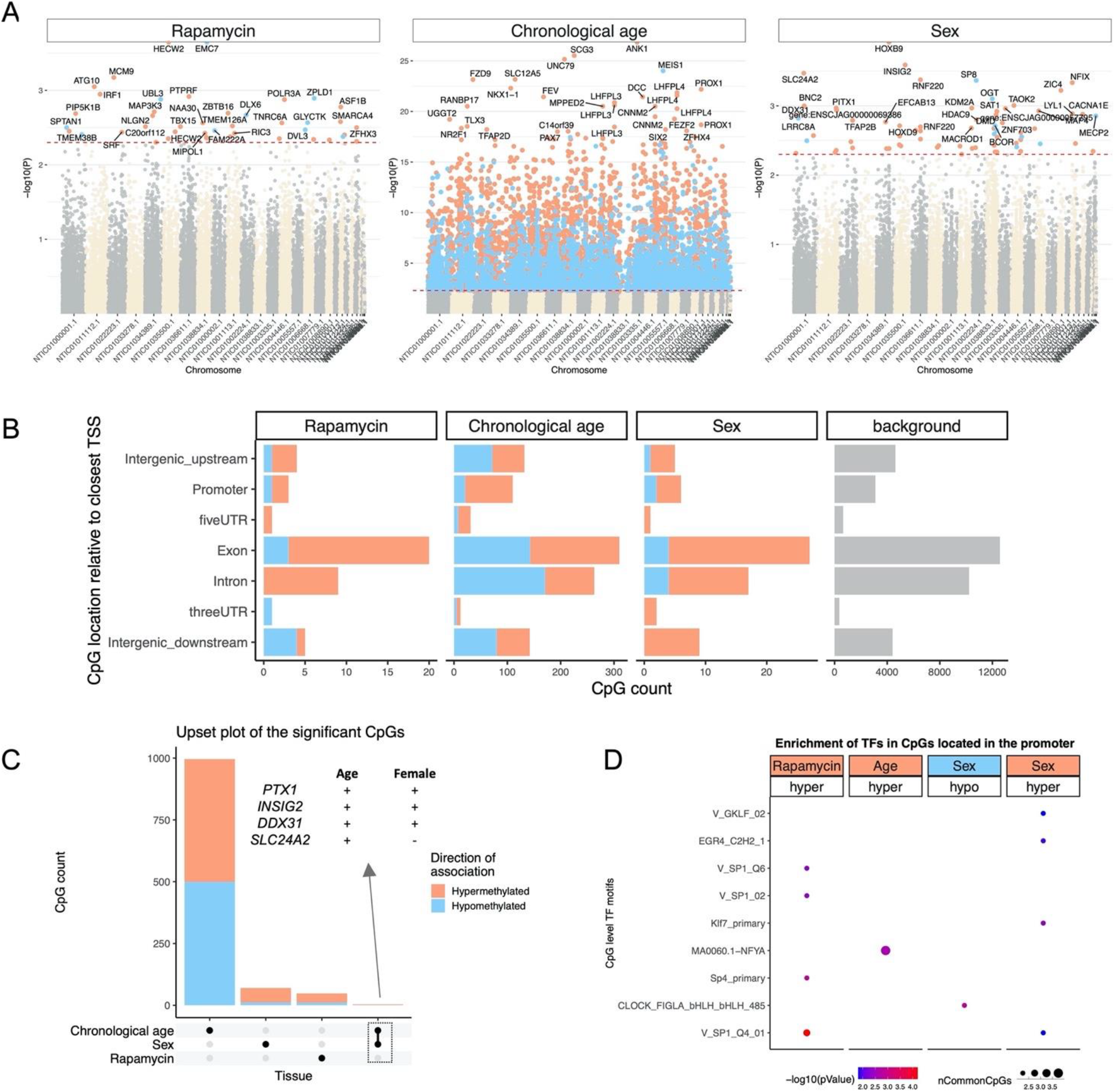
Epigenome-wide association (EWAS) of Rapamycin treatment, chronological age, and basal sex difference in blood of common marmosets (Callithrix jacchus). A) Manhattan plots of the EWAS of Rapamycin, chronological age, and sex. The rapamycin effect (N: Ct = 20, Rapamycin= 17) and Sex difference (N: M = 11, F = 9) was studied by multivariate regression model with chronological age as a co-variate. Animals with Rapamycin treatment were excluded for studying sex effects. The coordinates are estimated based on the alignment of Mammalian array probes to ASM275486v1.100 genome assembly. The direction of associations with p < 0.005 (red dotted line) is highlighted by red (hypermethylated) and blue (hypomethylated) colors. Top 30 CpGs was labeled by the neighboring genes. B) Location of top CpGs in each tissue relative to the closest transcriptional start site. Top CpGs were selected at p < 0.005 and further filtering based on z score of association with chronological age for up to 500 in a positive or negative direction. The number of selected CpGs: Rapamycin, 48; Age, 1000; and sex, 74. The grey color in the last panel represents the location of 35815 mammalian BeadChip array probes mapped to ASM275486v1.100 genome. C) Upset plot representing the overlap of aging-associated CpGs based on meta-analysis or individual species. The Neighboring genes of the top overlapping CpGs were labeled in the figure. D) Transcriptional motif enrichment for the top CpGs in the promoter and 5`UTR of the neighboring genes. The motifs were predicted using the MEME motif discovery algorithm, and the enrichment was tested using a hypergeometric test.

### Blood methylation study of rapamycin

Rapamycin (an inhibitor of mechanistic target of rapamycin, mTOR) is a promising anti-aging intervention because it has been observed to extend lifespan in laboratory rodent models. As a bridge towards clinical translation, we previously developed a cohort of aging marmosets at UT Health San Antonio (UTHSA) in which animals are treated daily with rapamycin with a primary aim of addressing the effect of this intervention on marmoset lifespan^28^. We have leveraged the existence of this cohort to address what effect rapamycin intervention has on the epigenetic clock. At the time of blood draw, animals had been on daily rapamycin treatment for approximately 2-3.5 years.

After dosing with rapamycin, we found no significant effect on the DNAmAge in blood in multivariate linear regression model (**Table 2**), which suggests no effect of rapamycin in DNA methylation age in relatively healthy, middle-aged marmosets.

**Table 2.** Multivariate regression model for evaluating the treatment effect of rapamycin in the test data. The dependent variable (DNAmAge) is based on an epigenetic clock that was trained (developed) in the training data (Table 1). Rapamycin does not have a significant effect on the DNAmAge of blood tissue in this study.

| Outcome: DNAmAge, R-sq.=61% |  |  |  |
| --- | --- | --- | --- |
|  | Coef | Std. Error | P.value |
| Age | 0.631 | 8.89E-02 | 3.38E-8 |
| Female | 0.667 | 4.75E-01 | 0.169 |
| Rapamycin | -0.181 | 4.44E-01 | 0.686 |

### EWAS of rapamycin

We also carried out an epigenome wide association study (EWAS) of rapamycin to identify individual CpGs whose methylation levels are related to treatment status. None of the CpGs were significant after correcting for multiple comparisons. Even at a very permissive significance threshold (alpha=0.005), we only found 48 CpGs (11 hypomethylated, 37 hypermethylated) that exhibited methylation change (**Figure 4A**). The top affected DNAm changes included hypermethylation in *HECW2* and hypomethylation in *EMC7* exons. In general, rapamycin mainly caused hypermethylation in all genic regions (**Figure 4B**). The affected CpGs by rapamycin showed neither a strong correlation with age nor a sex difference (**Figure 4C**). The top transcriptional factor motif that showed rapamycin-mediated hypermethylation was SP1 (**Figure 4D**), which is involved in several aging-associated diseases including cancer ^29^, hypertension ^30^, atherosclerosis ^31^, Alzheimer’s ^32^, and Huntington diseases ^33^. SP1 is also a key regulator of mTORC1/P70S6K/S6 signaling pathway ^34^, and also has several binding motifs in the RAPTOR promoter ^35^.

### Sex effects

In our previous mammalian methylation studies, we have found that one can accurately predict sex on the basis of DNA methylation profiles. Surprisingly, a random forest predictor of sex based on the marmoset methylation data generated relatively low accuracy of 66%, which did not improve after application of a generalized linear regression model. A random assignment of sex would have led to an accuracy of roughly 50% in this data set. Our EWAS of sex corroborates these results. None of the CpGs were significant at a genome wide significance level. Only a few CpGs (62 hypermethylated, 12 hypomethylated in females) in marmoset blood samples showed a sex effect at a very permissive significance threshold of alpha=0.005 (**Figure 4A**). *HOXB9* exon, a CpG downstream of *INSIG2*, *SLC24A2*, and *RNF220* intron were the top regions with hypermethylation in females. Most of the CpGs with sex differences showed hypermethylation in females (**Figure 4B**). Few of these CpGs were also positively associated with age (**Figure 4C**). SP1 was the top TF motif associated with hypermethylation in females and was also affected by rapamycin (**Figure 4D**). Overall, we find that sex differences in DNAm in marmosets is not as strong as that observed for all other mammalian species tested thus far.

## Discussion

Epigenetic clocks for humans have found many biomedical applications including the measure of age in human clinical trials ^10,36^. This instigated development of similar clocks for mammals such as mice ^18–23^. While rodent models have obvious advantages, it can be challenging to translate findings from rodents to primates. NHP will play an indispensable role in preclinical work of anti-aging treatments, and development of suitable biomarkers of age promises to greatly reduce the costs and time needed for carrying out studies in NHP.

Our novel DNA methylation data from 37492 CpG probes represent the most comprehensive dataset thus far tested in common marmosets. Due to genetic differences, a critical step toward crossing the species barrier was the use of a mammalian DNA methylation array that profiled methylation of CpGs that are flanked by sequences that are highly conserved across numerous mammalian species. These data allowed us to construct highly-accurate DNA methylation-based age estimation tools for the common marmoset that apply to the entire life course (from birth to old age).

The human-marmoset clock for relative age demonstrates the feasibility of building epigenetic clocks for two species based on a single mathematical formula. The single formula of the human-marmoset clock that is equally applicable to both species effectively demonstrates that epigenetic aging mechanisms are highly conserved. The mathematical operation of generating a ratio (age divided by maximum lifespan) also generates a much more biologically meaningful value because it indicates the relative biological age and possibly fitness of the organism in relation to its own species.

We expect that the availability of these clocks will provide a significant boost to the attractiveness of the marmoset as biological model in aging research. The use of novel animal models for research is often hampered by the lack of available molecular tools due to species differences. Due to the relatively species-agnostic approach of our methylation arrays, this clock has potential benefits beyond marmoset-specific research as well and reveal several salient features with regards to the biology of aging. First, the marmoset pan-tissue clock re-affirms the implication of the human pan-tissue clock, which is that aging appears to be a coordinated biological process that is harmonized throughout the body. Second, the ability to combine these two pan-tissue clocks into a single human-marmoset pan-tissue clock attests to the high conservation of the aging process across two evolutionary distant species. This implies, albeit does not guarantee, that treatments that alter the epigenetic age of marmosets, as measured using the human-marmoset clock is likely to exert similar effects in humans. If validated, this would be a step change in aging research.

In this regard, it is of interest that we found relatively little effect of rapamycin, a drug known to extend lifespan in mice and invertebrates, on the epigenetic age in blood samples from marmosets. It is thus not clear if this lack of effect suggests that rapamycin has no effect in this cohort of marmoset, or if the design of our study was unable to capture such differences. The expectation that rapamycin would impede ageing is predicated on the assumption that ageing and longevity are inextricably linked. Although this assumption is understandably intuitive, it remains to be proven. It is theoretically possible that some interventions that extend life, may not delay or slow-down ageing ^37–39^. This intriguing possibility remains to be tested in primates, and if the marmoset cohort on rapamycin is eventually observed to have extended lifespan, this may be a possible indication. Another factor to consider is the possibility that rapamycin effect on epigenetic age may only become apparent later, at older age, when physical and physiological differences between the control and rapamycin cohorts are manifested. However, we believe this explanation is unlikely because the treated animals had a mean age of 10 at the time of the blood draw (Table 1).

It would be of interest to further explore other interventions or tissue types in order to clarify these questions. In this cohort of animals, our ongoing study will determine the effect of rapamycin on both lifespan as well as several markers of healthy aging. These data will help clarify the relationship between rapamycin, aging and epigenetic age in the marmoset.

It is indeed surprising that one could not build an estimator of sex based on DNA methylation profile of marmoset. This study is part of a consortium that generated DNA methylation data from over 160 mammalian species, from which DNA methylation-based predictors of sex were readily developed for all except for marmosets. For example, random forest-based predictors of sex led to nearly 100 percent accuracy in all other species tested (including primates). By contrast, the out-of-bag accuracy of a random forest predictor was only 66% for blood samples from marmosets. We hypothesize that this reflects that marmosets are hematopoietic chimeras. The litter mates exchange stem cells across the placental anastomoses during development. The full biological mechanisms and consequences remain unclear. Indeed, there is still a debate regarding which tissues are truly chimeric, instead of appearing so due to “contamination” with chimeric blood product. Questions also remain regarding how the cells from two (or three) distinct genomes function in the body without significant genomic conflict. Our data suggest that potential differences in X inactivation in marmosets may drive the relatively lack of measurable changes in DNA methylation profiles used in our assays.

## Materials and Methods

### Animal care and maintenance

All marmosets used in this research were housed at the Barshop Institute for Longevity and Aging Studies at UT Health San Antonio (UTHSA). The Institutional Animal Care and Use Committee (IACUC) of UTHSA is responsible for monitoring housing and animal condition regularly to ensure all guidelines are met for the safety and health of the animals. This research was reviewed and approved by the UTHSA IACUC and experiments were conducted in compliance with the US Public Health Service’s Policy on Humane Care and Use of Laboratory Animals and the Guide for the Care and Use of Laboratory Animals and adhered to the American Society of Primatologists (ASP) principles for the ethical treatment of non-human primates. Animals used in this study were based on age as well as a record of relatively good health as assessed by veterinary examination. Animals in this study were born either at UTHSCSA or transferred from the Southwest National Primate Research Center (SNPRC) in San Antonio TX. At UTHSA, animals were maintained using a modified specific pathogen-free barrier facility ^40^.

Each animal received three diet choices daily provided ad libitum: Harlan Teklad purified marmoset diet (TD99468), Mazuri Callitrichid gel diet (5MI5) and ZuPreem.

### Rapamycin

Rapamycin encapsulated in Eudragit (and Eudragit alone) were purchased from Rapamycin Holdings (San Antonio TX). Marmoset were trained to receive oral dosing of rapamycin (or eudragit alone) mixed into yogurt ^41^. Once trained, marmosets were treated once daily with a dose of rapamycin roughly equivalent to 1 mg rapamycin/kg body weight. Animals were dosed Monday-Friday with dose received between 08:00 and 10:00 but not dosed on Saturday or Sunday.

Blood draws and clinical blood counts/chemistry: All blood draws were taken during the morning hours of 08:00-11:00 from fed, non-anesthetized animals restrained in a custom assembly. Femoral vein blood collection (1.0-2.0 mL) was performed on each animal and blood was placed in to PAXgene blood collection tubes (Qiagen). Whole blood was frozen and stored at −80° C for shipment to UCLA for methylation analysis.

#### Study samples

For these studies we selected samples representing the entire primate lifespan, from neonate to old age. Genomic DNA was isolated from tissue samples mostly using Puregene chemistry (Qiagen). DNA from liver was extracted manually and from blood using an automated Autopure LS system (Qiagen). From old liver tissues and clotted blood samples DNA was extracted manually using QIAamp DNA Blood Midi Kit and the DNeasy Tissue Kit according to manufacturer’s protocol (Qiagen, Valencia, CA). DNA from BA10 was extracted on an automated nucleic acid extraction platform Anaprep (Biochain) using a magnetic bead-based extraction method and Tissue DNA Extraction Kit (AnaPrep).

#### Rhesus monkey

To validate the vervet clock cross species, we utilized n=240 methylation arrays profiling blood, cortex, and liver samples described in a companion paper ^25^. The methylation data were generated on the same array platform (MammalMethylChip40).

#### Vervet monkey

To validate the vervet clock cross species, we utilized n=240 methylation arrays profiling blood, prefrontal brain cortex and liver from the vervet monkey described in a companion paper ^24^. The methylation data were generated on the same array platform (MammalMethylChip40).

#### Human tissue samples

To build the human-marmoset clock, we analyzed previously generated methylation data from n=850 human tissue samples (adipose, blood, bone marrow, dermis, epidermis, heart, keratinocytes, fibroblasts, kidney, liver, lung, lymph node, muscle, pituitary, skin, spleen) from individuals whose ages ranged from 0 to 93. The tissue samples came from three sources. Tissue and organ samples from the National NeuroAIDS Tissue Consortium ^42^. Blood samples from the Cape Town Adolescent Antiretroviral Cohort study ^43^. Skin and other primary cells provided by Kenneth Raj ^44^. Ethics approval (IRB#15-001454, IRB#16-000471, IRB#18-000315, IRB#16-002028).

#### DNA methylation array

All DNAm data used was generated using the custom Illumina chip “HorvathMammalMethylChip40”, so called the mammalian methylation array. The mammalian methylation array is an attractive tool for DNAm assessment, because it comprises 37492 probes, including nearly ~36 K probes targeting CpG sites in highly conserved regions in mammals. 1951 probes were selected based on their utility for human biomarker studies: these CpGs, which were previously implemented in human Illumina Infinium arrays (EPIC, 450K) were selected due to their relevance for estimating age, blood cell counts, or the proportion of neurons in brain tissue. The remaining probes were chosen to assess cytosine DNA methylation levels in mammalian species (Arneson, Ernst, Horvath, in preparation). The particular subset of species for each probe is provided in the chip manifest file can be found at Gene Expression Omnibus (GEO) at NCBI as platform GPL28271. The SeSaMe normalization method was used to define beta values for each probe ^45^.

### Penalized Regression models

Details on the clocks (CpGs, genome coordinates) and R software code are provided in the Supplement. Penalized regression models were created with glmnet ^46^. We investigated models produced by both “elastic net” regression (alpha=0.5). The optimal penalty parameters in all cases were determined automatically by using a 10 fold internal cross-validation (cv.glmnet) on the training set. By definition, the alpha value for the elastic net regression was set to 0.5 (midpoint between Ridge and Lasso type regression) and was not optimized for model performance.

We performed a cross-validation scheme for arriving at unbiased (or at least less biased) estimates of the accuracy of the different DNAm based age estimators. One type consisted of leaving out a single sample (LOOCV) from the regression, predicting an age for that sample, and iterating over all samples. A critical step is the transformation of chronological age (the dependent variable). While no transformation was used for the blood clock for marmosets, we did use a log linear transformation for the dual species clock of chronological age (Methods).

### Relative age estimation

To introduce biological meaning into age estimates of marmosets and humans that have very different lifespan; as well as to overcome the inevitable skewing due to unequal distribution of data points from marmosets and humans across age range, relative age estimation was made using the formula: Relative age= Age/maxLifespan where the maximum lifespan for the two species was chosen from the *anAge* data base ^47^. The maxLifespan of marmosets and humans was chosen as 22.8 and 122.5, respectively.

### Epigenome wide association studies of age

EWAS was performed in each tissue separately using the R function “standardScreeningNumericTrait” from the “WGCNA” R package^48^. Next the results were combined across tissues using Stouffer’s meta analysis method.

## Acknowledgements

This work was supported by the Paul G. Allen Frontiers Group (SH). National Institute of Aging 1U19AG057758, R01 AG050797, P30 AG013319, P30 AG044271

ABS was also supported Geriatric Research, Education and Clinical Center of the South Texas Veterans Health Care System. This material is the result of work supported with resources and the use of facilities at South Texas Veterans Health Care System, San Antonio, Texas. The contents do not represent the views of the U.S. Department of Veterans Affairs or the United States Government.

## SUPPLEMENTARY MATERIAL

**Supplementary Figure 1.**
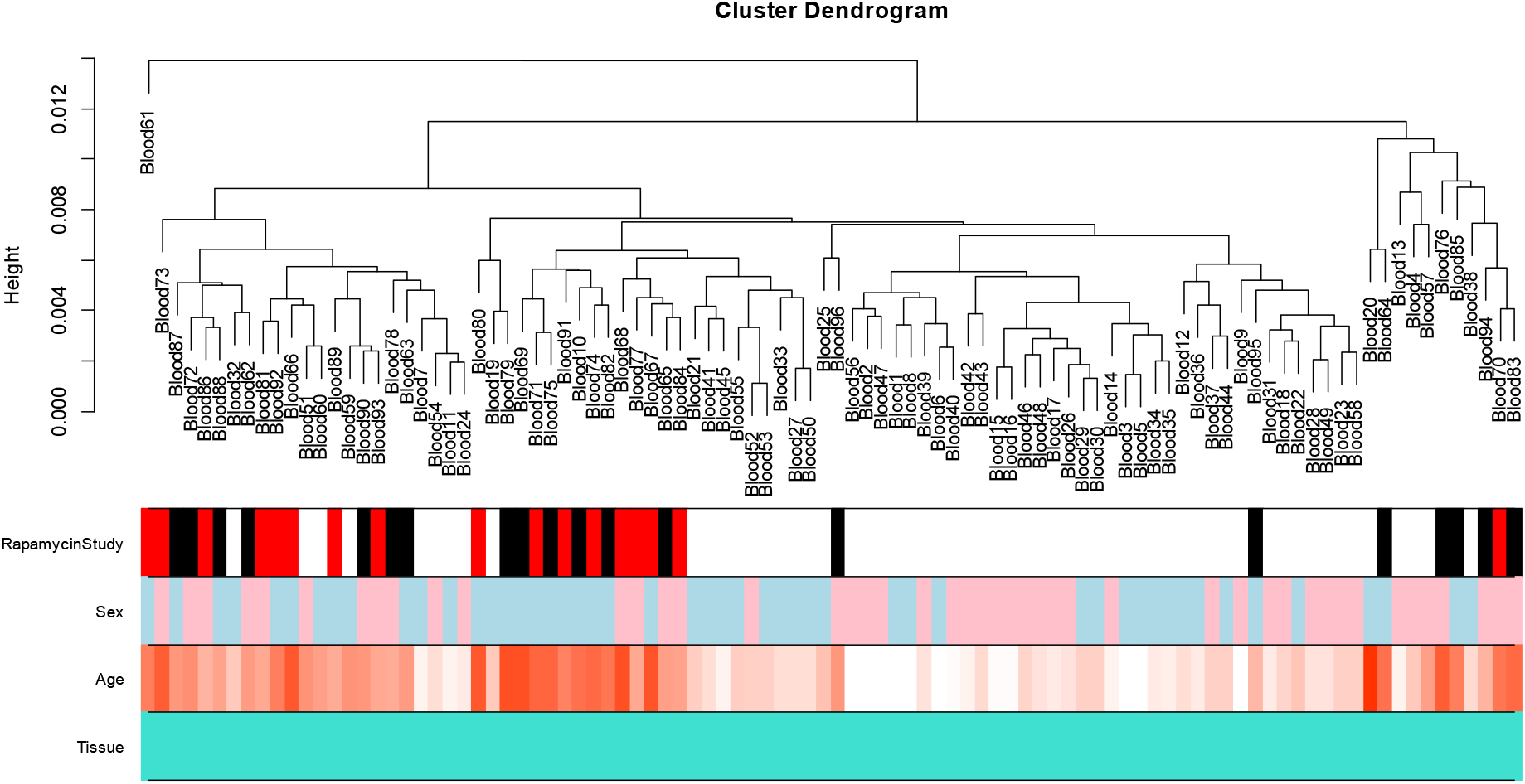
Unsupervised hierarchical clustering of blood samples from marmosets. Average linkage hierarchical clustering based on the inter-array correlation coefficient (Pearson correlation). The low height values (y-axis) indicate high inter array correlations (R>0.98) and high quality. There is no obvious clustering pattern. The first color band indicates rapamycin treatment status: red=rapamycin treated, black=control, while=not part of the rapamycin study.

